# Direct evidence of acid-driven protein desolvation

**DOI:** 10.1101/2025.09.01.673474

**Authors:** Farzad Hamdi, Ioannis Skalidis, Inken Kaja Schwerin, Jaydeep Belapure, Dmitry A. Semchonok, Fotis L. Kyrilis, Christian Tüting, Johannes Müller, Georg Künze, Panagiotis L. Kastritis

**Affiliations:** Department of Integrative Structural Biochemistry, Institute of Biochemistry and Biotechnology, Martin Luther University Halle-Wittenberg, Kurt-Mothes-Straße 3, 06120 Halle/Saale, Germany; Interdisciplinary Research Center HALOmem, Charles Tanford Protein Center, Martin Luther University Halle-Wittenberg, Kurt-Mothes-Straße 3a, 06120 Halle/Saale, Germany; Structural Biochemistry, Bijvoet Centre for Biomolecular Research, Utrecht University, 3584 CG Utrecht, the Netherlands; Institute for Drug Discovery, Leipzig University, Brüderstraße 34, 04103 Leipzig, Germany; Navigo Proteins GmbH, 06120, Halle/Saale, Germany; Instituto de Tecnologia Química e Biológica António Xavier, Universidade Nova de Lisboa. Av. da República, 2780-157 Oeiras, Portugal; Institute of Chemical Biology, National Hellenic Research Foundation, 11635 Athens, Greece; Institute of Pharmacy, Martin-Luther-University Halle-Wittenberg, Weinbergweg 22, D- 06120 Halle (Saale), Germany; Interdisciplinary Center for Bioinformatics, Leipzig University, 04107 Leipzig, Germany; Center for Scalable Data Analytics and Artificial Intelligence, Leipzig University, 04105 Leipzig, Germany; Biozentrum, Martin Luther University Halle-Wittenberg, Weinbergweg 22, 06120 Halle/Saale, Germany

## Abstract

Water and its ability to modulate the protonation states of biomolecules govern the physical chemistry of life, dictating their metabolic functions(*1*). However, how amino acid protonation alters protein hydration and solubility(*2, 3*) is an open question since Kuntz and Kauzmann proposed p*H*-driven protein desolvation in 1974(*4*). Here, in a series of high-resolution cryo-electron microscopy structures of a protein complex at different p*H* values (from p*H* 9.0 to 3.5) we examined thousands of observable hydration sites. Cryo-EM data, in agreement with constant-p*H* molecular dynamics simulations, show that nearly half of protein-bound waters exchanged with the bulk solvent upon acidification, with ∼100 waters lost per p*H* unit per molecule. The loss of waters was most significant around the side chains of glutamate and aspartate residues while specific polar residues, mostly asparagine, anchored persistent waters. A positionally conserved hydration layer was observed across all p*H* conditions, accounting for 40% of resolved waters. Those waters displayed denser packing than less persistent waters, forming a p*H*-independent solvation shell. Acid-induced water exchange also displaced bound iron, providing a mechanistic link between solvation and metal release. Our findings demonstrate the core principles of acid-driven protein desolvation, resolving a 50-year-old biochemical hypothesis(*4*).

## Main

Water molecules produce complex solvation layers around macromolecules that may extend beyond 20 Å(*5*). Their dynamics correlate with protein structure, function, interaction and dynamics(*1, 2, 6*). Atomic modelling of ordered water molecules in high-resolution 3D electron density maps(*7*), revealed their roles in critical biological processes, such as RNA function(*8*), peptide bond cleavage(*9*), signal transduction by GPCRs(*10*), e.g., in rhodopsin, and proton-translocation pathways in the respiratory pathway(*10*) or in photosynthesis(*11*). Such water molecules, typically resolved in high-resolution structures, are classified as “structural” or “ordered waters”, *i.e*., integral parts of the macromolecular architecture(*6*) and together with faster, low-occupancy waters define solvation dynamics(*12*). X-ray crystallography(*13*) and nuclear magnetic resonance(*14*) (NMR) have been traditionally applied to experimentally probe the structure and dynamics of water molecules, respectively, while AI-based approaches do not, yet, model water positions in biomolecular interactions at near-atomic resolution(*15*) or, when they do, response to changes in ionic strength is not considered(*16*). The dynamics of the water shell that surrounds proteins have been studied in great detail by methods such as NMR, terahertz time-domain spectroscopy and neutron scattering, amongst others, to reveal that water molecules around proteins are much more ordered in comparison to bulk water(*12, 17-25*). Interestingly, NMR studies employing, e.g., nuclear Overhauser effect spectroscopy (NOESY) have shown that there is no difference in the residence time of surface water molecules which are observed in X-ray structures versus those that are not detected(*26*).

Single-particle cryo-electron microscopy (cryo-EM) is an alternative method for determining water molecule structures interacting with biomolecules(*27*) - but its application has been limited, being the most recent of those in terms of high-resolution analysis. Cryo-EM structures where water molecules can be resolved have recently become available due to the ongoing resolution revolution(*28*), following advances in instrumentation, image processing and analysis software, and integration with standard protein modelling software(*29*). An intrinsic advantage of cryo-EM is its ability to probe macromolecular structures of proteins that are rapidly frozen in vitreous ice preserving near-native conformations without requiring crystallization(*30*). Therefore, specimens can be compared under different physical-chemical conditions to probe their structural response, e.g., the temperature dependence of enzyme catalysis(*31*). This is particularly important when studying the solvation layer and its response to the concentration of hydronium ions in aqueous macromolecular solutions. Indeed, p*H* dependency of proteins has been well known to have critical functional implications(*32*), driving conformational changes(*33*), dynamics(*34, 35*) and, ultimately, cellular function(*36*). At the molecular level, p*H* can affect the hydration layers of macromolecules(*4*), and importantly, local pKa of crucial residues due to localized protonation changes. Minuscule changes might render an atom from a nucleophile to an electrophile, and thereby, allowing certain biochemical reactions to occur. Kunz and Kauzmann(*4*) observed via ultracentrifugation, slight density increases in protein sedimentation experiments as a function of decreasing p*H* and interpreted them as changes in the amount of bound water resulting from changes in the ionization state of protein carboxyl groups.

Despite decades of research on the effect of p*H* on bound water with an array of biophysical(*37-39*) and biocomputing(*40*) methods, experimental observation of individual water molecules and their dependence on p*H* has yet to be probed. Here, we have undertaken this task and analyzed waters in the structure of human apoferritin - a biomolecule that can withstand such drastic changes in p*H*(*41*), at varying hydronium ion concentrations. Apoferritin was selected as an example because of its highly stable nature(*42*), making it an ideal specimen for cryo-EM(*43, 44*), previous experience of the laboratory in its high-resolution structure determination(*45*), and excellent tolerance to a wide range of p*H*(*41*).

## Results

### Recovery of high-resolution structures at pH 3.5-9.0

We overexpressed the human apoferritin gene construct and purified the protein complex (**Methods, Fig. S1A-B**), recovering a monodisperse peak of the molecule in size exclusion chromatography at an expected molecular weight of 450 kDa (**Fig. S1C**). Before vitrification, we optimized the p*H* of the retrieved aqueous solution using a broad range buffer series(*46*). The selected range ensured that the structural effects to be observed are not affected by any distinct composition of the aqueous solution studied and are effects of adjusted p*H* solution. Cryo-electron microscopy (cryo-EM) screening showed that the molecule was stable in the p*H* range of 3.5-9.0 (**Fig. S2A**), as, at p*H* ∼3, the molecule dissociates(*47*) **(Fig. S2B-D)**. After data collection (**Table S1**), image processing (**Fig. S3**), and subsequent refinement procedures (**Table S1, Methods**), we were able to reconstruct cryo-EM maps of human apoferritin at p*H* 3.5, 4.0, 5.0, 7.0, and 9.0 at resolutions around 2 Å (**Table S1, Fig. S3**). Water molecule densities in all modelled structures were highly visible while accurate modelling of water molecules, co-factors, and side chains (**Fig. S4**) was feasible at this resolution.

Cryo-EM maps did not show meaningful p*H*-dependent size deviations as previously reported by atomic force microscopy(*48*), in agreement with low-resolution small angle X-ray scattering data(*41*). Instead, at varying p*H* conditions, apoferritin suspended in vitreous ice retains its expected molecular structure and higher order structure. Specifically, the protein backbone remained stable, with Cα root-mean-square-deviation (RMSD) values relative to p*H* 7 being small, *i.e*., 0.75 Å at p*H* 3.5, 0.49 Å at p*H* 4, 0.62 Å at p*H* 5, and 0.32 Å at p*H* 9, corroborating that changes in non-polypeptide components are not due to large-scale protein rearrangements. To investigate the structural stability and dynamics of apoferritin at different p*H* values at the atomic level, we conducted all-atom molecular dynamics (MD) simulations. We simulated the varying p*H* conditions by adjusting the charge state of protonatable residues (Glu, Asp, Lys, His) according to the pKa values of their side chain functional groups, which we determined using p*H* titration calculations (see **Methods, Fig. S5**). In agreement with the cryo-EM data, the apoferritin complex remained close to the starting structure in the MD simulations conducted at p*H* values of 5.0 and above. However, with the side chain protonations adjusted to mimic p*H* values of 3.5 and 4, the structure deviated more strongly from the starting coordinates (backbone RMSD of ∼3 Å, **Fig. S6**), indicative of an increased instability of the apoferritin complex at low p*H* values. This structural deviation at lower p*H* likely reflects residues with pKa values slightly above 4. In our simulations, these residues were modeled as fully protonated, leading to changed protein solvation and intermolecular interactions, whereas under the cryo-EM conditions, these residues may remain partially deprotonated or constitute a minority of the apoferritin molecules that did not end up in the final reconstruction during processing.

### Distinct pH-dependency of bound water molecules and metals

The modelled structures within the high-resolution cryo-EM maps provided us with a series of hypotheses. First, we wanted to investigate if modelled water molecules exhibited conformational variation higher than that of amino acid residues or bound metals, as it is known from NMR systematic investigations that water reorientation and translation at the protein surface is 3-5 times slower than in bulk(*6*). To address this, we estimated the per-atom resolvability in all five cryo-EM maps. Results show that the resolvability of atoms, calculated by their Q-scores and translated into B-factors(*49*), is comparable for all distinct chemical species, irrespective of the p*H* in which apoferritin was buffered (Q-scores > 0.8, **Fig. 1A**). This exercise corroborates the observation that resolvability and confidence for the positions of water molecules are not compromised as a function of p*H* or slight differences (∼0.1 Å) around 2 Å resolution.

**Figure 1.**
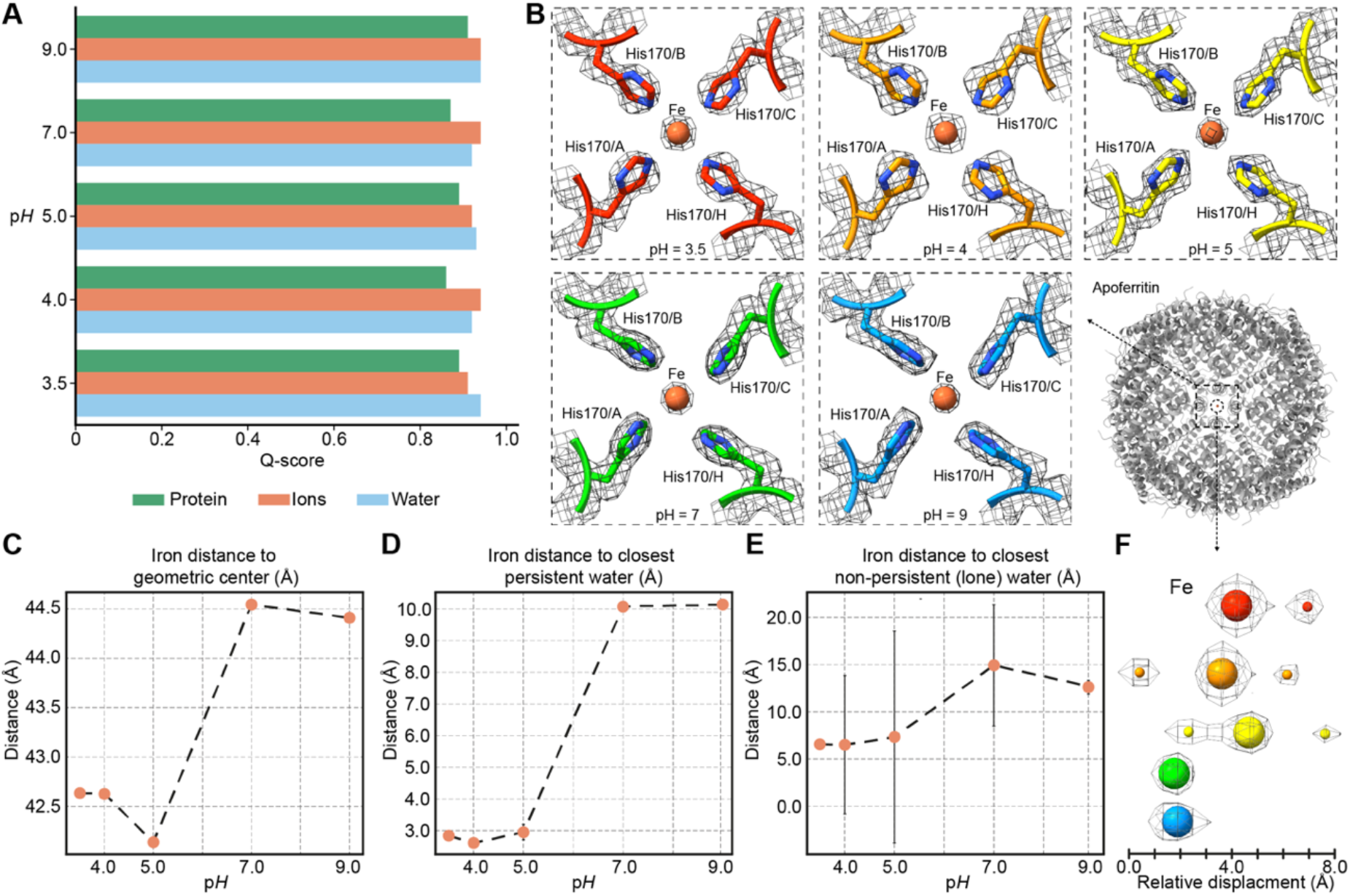
Analysis of atom resolvability and iron positioning in apoferritin cryo-EM maps determined at varying p*H*. (**A**) High Q-scores are calculated for the resolved Coulomb potential maps corresponding to residues (green), bound ions (brown) and water molecules, (blue) at all p*H* values. (**B**) The iron is resolved in all cryo-EM maps, coordinated by histidines which show minimal side-chain conformational changes. (**C**) Calculation of distances from the apoferritin geometric center to individual iron ions in cryo-EM reconstructions. (**D**) Distances of persistent and **(E)** non-persistent (lone) water molecules in proximity to the iron ion. Persistent correspond to water molecules always present in all p*H* values, non-persistent are not. Only persistent water molecules are identified in proximity to the iron, with their distances getting shorter with decreasing p*H* (at pH 5 and below, distances correspond to those of a hydrogen bond). (**F**) Relative positional change of the bound iron in the channel sites. Proximal water molecules appear at p*H* values of 5 and lower.

Next, we studied the iron ion coordination in the resolved structures for the different p*H* conditions. Apoferritin is a ferroxidase that converts Fe(II) to Fe(III), and internalizes the iron in its core; however, the transfer of iron at the atomic level is not well understood(*50*). Previously resolved structures show a network of histidines organizing the iron(*51*). In our structures, we have unambiguously determined an iron ion at the iron entry channels, corresponding to 8 bound iron ions in total (**Fig. 1B**). Because we resolved the molecule at different p*H* values, we sought to directly observe the localization of the iron and its coordination as a function of decreasing p*H*. This may reflect the general principle that metal solubility increases at lower p*H*, as protons compete with metal ions for coordination sites. It is also critically connected to cellular iron homeostasis(*52*), and overall, important for iron homeostasis, *e.g*., in leukamias(*53*) or neurodegeneration(*54*).

We observe that with decreasing p*H*, the iron is displaced towards the geometric center of the entire protein complex (**Fig. 1C-F**). This behavior is highly specific for the iron ion and not for the structural magnesium (**Fig. S7A-C**). This discrete behavior of bound metals as a function of p*H* underlines the fine-tuning of metal local coordination and further exchange with the bulk solvent. We then specifically asked if these metals were also further coordinated by stable water molecules in their proximity. Stable water molecules are often identified in metal-binding sites of proteins and are directly related to function(*55*). Structural water is present in proximity to the iron (**Fig. 1D**) but does not persist with increasing p*H*. Whereas at p*H*=5.0 or lower, a water molecule appears in the proximity of the iron ion, coordinated by four histidine residues and further stabilized by hydrophobic interactions with nearby leucines, this interaction disappears at a higher p*H* (**Fig. 1D**). Therefore, we identified at least one example in the high-resolution apoferritin structure where p*H* effects can directly drive the solvation of its iron channel site combined with a correlated inward movement of the iron (**Fig. 1C, 1F**) mediated by acid-forming water-metal hydrogen bonds (**Fig. 1D, Fig. S8**).

### pH-dependent protein desolvation

Kuntz and Kauzmann predicted, based on sedimentation data at different p*H*, that biomolecules are desolvating with decreasing p*H*(*4*). Due to the high-resolution structures that we determined at different p*H*, we can validate or reject this prediction. After modelling bound water molecules in all five cryo-EM maps (**Fig. S4, Table S1**) and unambiguously showing comparable resolvability with amino acid residues and metals (**Fig. 1A**), we counted the bound water molecules in the complete 24-mer structures (**Table S1)**. An apparent decrease in the number of bound water molecules upon acidification can be observed (**Fig. 2A**). A trend is revealed where ∼100 bound water molecules are released per *pH* unit (R≈0.97). This trend can be well reproduced when analyzing the number of water binding sites in the corresponding MD simulations (**Fig. 2A**). When the p*H* is lowered from 9 to 3.5, the number of resolved water molecules within a distance of 5 Å from the protein surface decreased by almost 50%.

**Figure 2.**
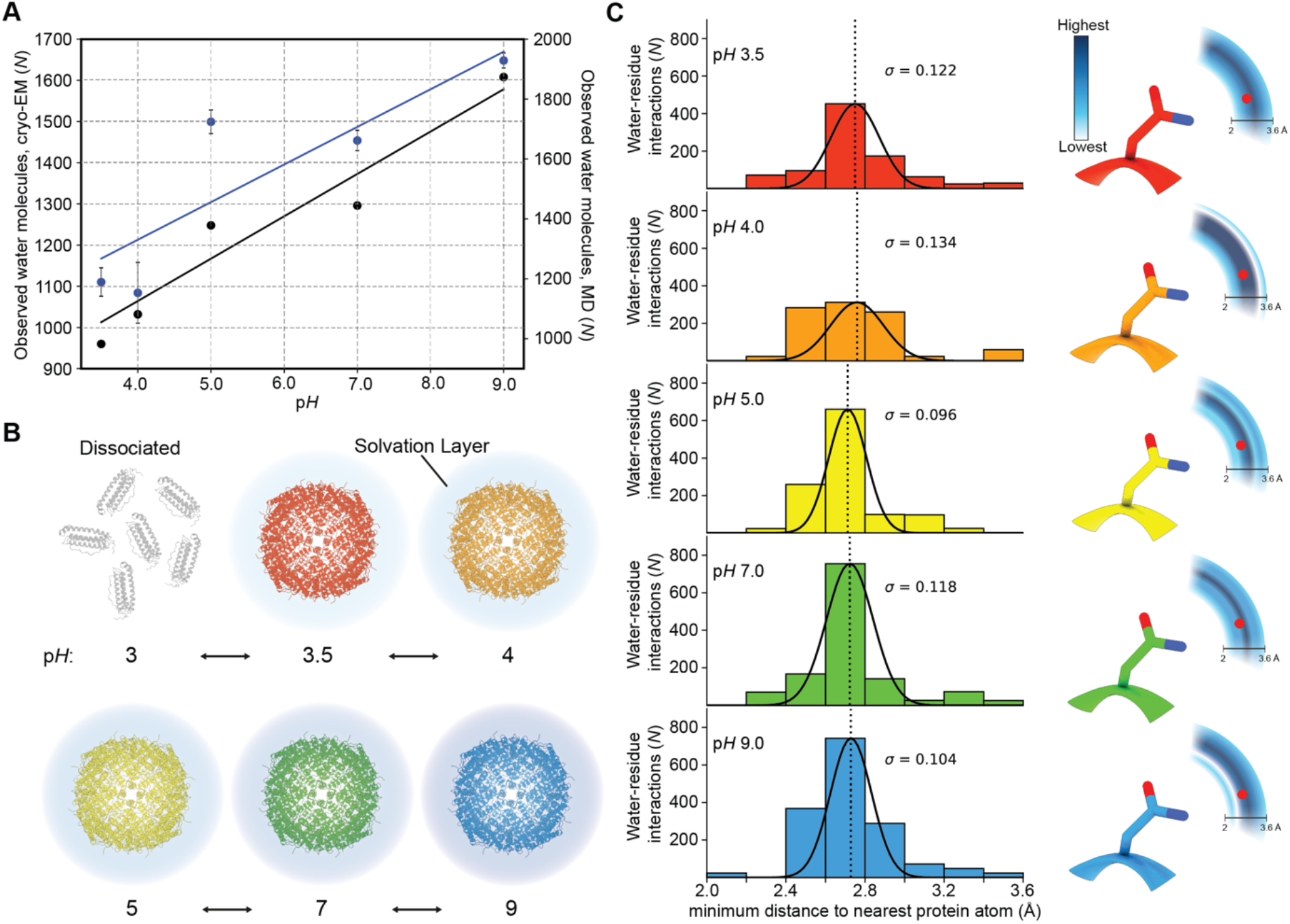
Solvation layer density and localization across p*H*. (**A**) Total number of surface-bound water molecules determined from the cryo-EM maps (black points) or by running MD simulation (blue points), respectively, as a function of *pH*. A clear upward trend is visible with increasing p*H*. (**B**) Graphical representation of solvation layer density. At p*H* 3, there is no complex formation, whereas, with increasing p*H*, the density of the visible water solvation layer increases. (**C**) Distribution of distances of resolved water molecules to the nearest amino acid atom. There is a clear Gaussian distribution in all p*H*s with upper bounds at 3.6 Å, lower bounds at 2 Å and an average of ∼2.7 Å. On the right, a 2D displacement diagram illustrates a water molecule adjacent to a protein residue, interpreting the distance distributions shown on the left.

This unprecedented observation can effectively rationalize the origins of acidic p*H*-induced dissociation at p*H*<3.5 (**Fig. 2B**). Effectively, the exchange of bound water to bulk is gradually increased up to a threshold; above this threshold, at p*H*<3.5, solvation of the 24-mer can no longer be stabilized once the water molecules nearest to its surface become too disordered to maintain effective solvation, eventually dissociating the higher-order structure to its more stable monomers (model in **Fig. 2B**). Next, we asked if the resolved bound water molecules exhibit similar distance deviations as those determined for the iron ion (**Fig. 1B**). Results show that distances of water molecules follow a complex behavior: At low p*H*, they have a less canonical distance to their nearby atoms as shown by the flattening of their distance distribution (**Fig. 2C**). In contrast, in higher p*H*, equivalent distribution is tighter (**Fig. 2C**) consistent with faster water motions as observed before(*56*). This observation also aligns with Kuntz and Kautzmann(*4*) who hypothesized that protein hydration reflects an ensemble of bound water states with varying mobilities rather than a single homogeneous population. It is also important to take into account changes in protein flexibility as a function of p*H*, which could affect or even orchestrate protein hydration. Even if we do not observe major conformational changes of apoferritin across p*H*, minor populations might still exist, as also shown by the MD simulations at acidic p*H*. Furthermore, studies on human serum albumin that populates at least five states as a function of p*H* (ranging from an extended state at p*H* 2.7 to an ‘aged’ state at p*H* 10) showed discrete hydration dynamics for each state(*56*), in agreement with our observation **(Fig. 2C)**. Our study establishes that acidic p*H*-dependent protein dissociation is coupled to progressive water layer flexibility and redistribution of bound water molecules, hydration being the key determinant of structural stability.

### Side-chain specific contributions to bound water molecule retention

Protein solvation is influenced by global p*H* and residue-specific protonation states(*57*). We therefore asked whether the observed changes in hydration arise from p*H*-dependent shifts in residue charge. To investigate if the p*H*-dependent hydration differences in the structure are linked to specific residue types, we calculated per-residue bound water molecule counts as a function of p*H* and compared these values with the per-residue hydration site numbers obtained from the MD simulations. This analysis showed that certain residue types get considerably desolvated with decreasing p*H* while others show only a small or no significant decrease of their number of bound water molecules (**Fig. 3A**). Specifically, glutamate and aspartate side chains have the largest number of bound waters of all amino acids at p*H* 9 and show the most pronounced desolvation with decreasing p*H*. Interestingly, this effect is also observed for basic residues (lysine and arginine) and to a lesser extent for histidine and polar residues (glutamine, asparagine, threonine, serine, tyrosine) (**Fig. 3A**). As expected, residues with apolar side chains (alanine, valine, leucine, isoleucine, phenylalanine, tryptophan, methionine, proline) or glycine show the smallest bound water numbers and least p*H*-dependent desolvation. Although low-resolution methods such as X-ray photo-electron spectroscopy have previously described p*H*-dependent desolvation effects and related them to changes in the residues’ charge state and hydrogen bonding capability with water(*58*), we here directly visualize and quantify this effect at an atomic level.

**Figure 3.**
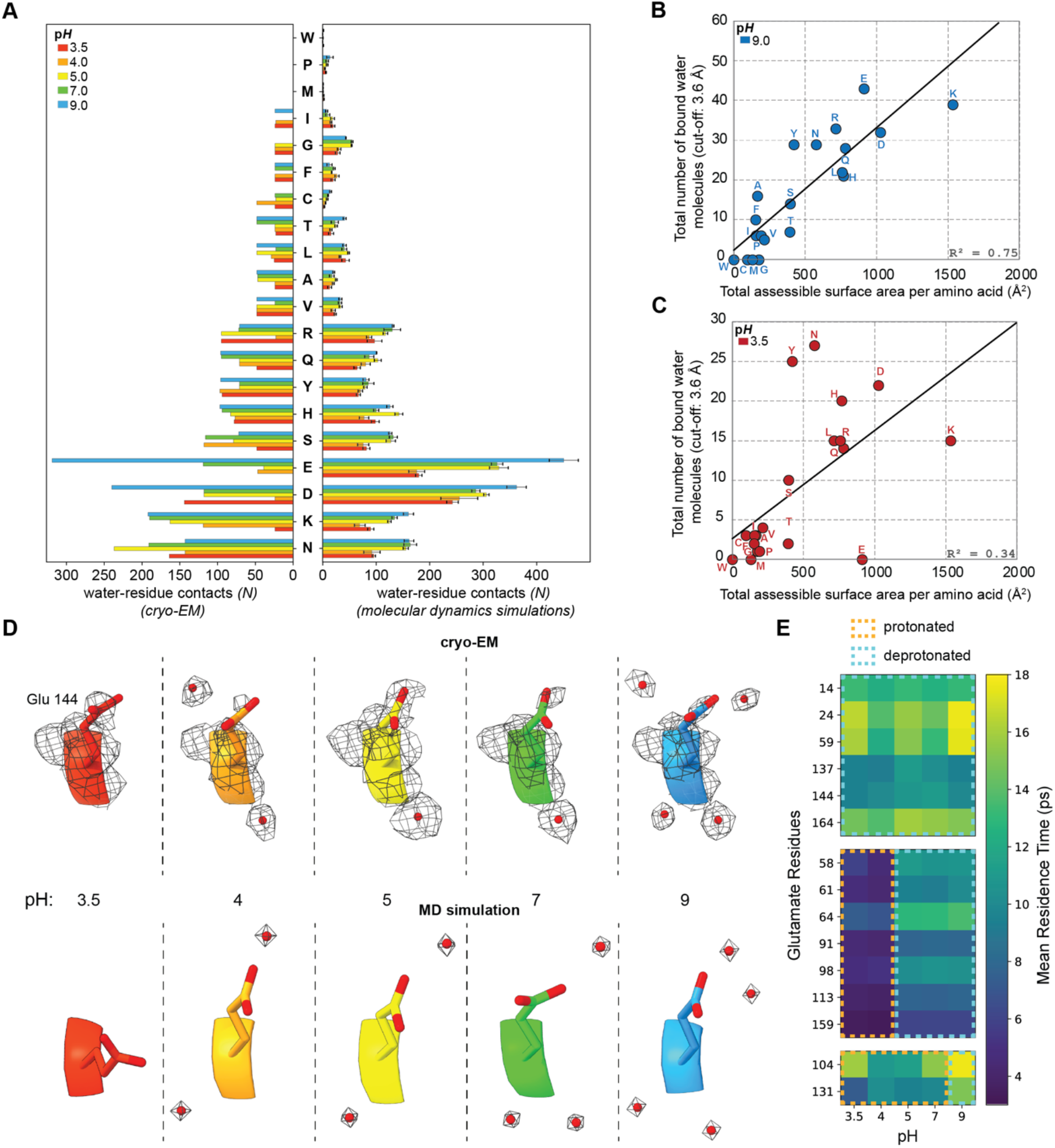
Per-residue bound water molecule counts at different p*H* values. (**A**) Total modelled water count per p*H* in proximity to each amino acid type in the cryo-EM maps (left) and in the MD simulations (right). **(B-C)** p*H*-dependent relationship between amino acid surface area and bound water molecules in apoferritin. Amino acid-specific counts of bound water molecules within 3.6 Å are plotted against their total accessible surface area, for each residue. Each point corresponds to all residues of a given amino acid type within the apoferritin structure. Correlations are strong, but become increasingly weaker at increasingly lower p*H* **(Fig. S9)**. Acidic and charged residues display a decreased water count in their proximity compared to other residue types. **(D-E)** Residence times and positions of waters surrounding glutamates. **(D)** Visualization of Glu144 and its surrounding water molecules across p*H* in the cryo-EM maps and MD simulations. **(E)** Residence times of waters surrounding the glutamate residues at different simulated p*H* values.

To evaluate if p*H*-dependent desolvation effects observed are generalizable, we correlated identified bound waters with total accessible surface area **(Fig. 3B-C, Fig. S9A)** and hydrophilicity (**Fig. S9B**) for each amino acid, deriving strong correlations at neutral and alkaline p*H* (R^2^ up to 0.75) **(Fig. 3B)**. At these conditions, overall hydration is largely dictated by surface area and polarity **(Fig. S9B)**, as expected(*59, 60*). However, at lower p*H*, correlations weakened markedly (R^2^ as low as 0.24), reflecting a gradual breakdown of residue-specific hydration **(Fig. 3C, S9A)**. Acidic side chains lost most bound waters upon protonation, whereas polar residues retained waters. These results show that protein hydration is organized and residue type-dependent at high or neutral p*H*, but collapses under increasingly acidic conditions, where many waters are displaced to bulk solvent.

To investigate how the persistency of water molecules in the macromolecular structure is affected by p*H* changes, we calculated the occupancy and residence time for the water binding sites that interacted with the amino acid residues in the MD simulations. The most considerable changes in the water molecule persistence were observed for glutamate residues, which showed a progressive reduction in their bound water molecule number with decreasing p*H*. For example, **Fig. 3D** illustrates the water sites interacting with residue E144, with the water positions closely matching between cryo-EM structure and the MD simulation. While at p*H* 9, E144 is interacting with 4 water molecules, it is bound to only 2 at p*H* 4.0 and is entirely desolvated at p*H* 3.5. The water residence times of glutamates also vary substantially from 4 ps up to 18 ps (**Fig. 3E**). We noted three groups of glutamate residues, demonstrating distinct water stabilities. The first group of glutamates has low pKa values of 2 to 4, as determined by constant p*H* MD simulations (see **Methods, Fig. S5**), and are deprotonated at all p*H* values. These residues show high residence times of 10 ps or higher, which vary minimally between different p*H* levels, reflecting stable water binding. The second group of glutamates has pKa values between 4 and 5 (**Fig. S5**). Consequently, these residues are protonated below p*H* 5 and deprotonated when the p*H* is 5 or higher. The change in the side-chain protonation state of these glutamates is accompanied by a jump of their bound water residence times from 4 ps at p*H* ≤ 4 to around 10 ps at p*H* ≥ 5, possibly due to stronger hydrogen bonding of their associated water molecules when the side chains get deprotonated. The third group includes two glutamate residues (E104, E131), which have exceptionally high pKa values (pKa_E104_ = 8.9 and pKa_E131_ = 7.9) and exist in a protonated state at p*H* ≤ 7. Their side-chain deprotonation at p*H* 9 leads to increased water residence times, reflecting again more stable water binding. In addition to glutamates, similar trends in the dynamics of bound water molecules, due to changes in the residues’ pKa values and protonation states, can be observed also for aspartates (**Fig. S10)**. Specific structural water molecules occur close to specific amino acids, as shown in **Fig. 3**.

Finally, to understand global retention propensities of bound water across p*H* values, we clustered water oxygens based on their distance (1.0 Å cutoff) across aligned modelled structures along with a nearest-neighbor residue assignment using a 3.6 Å cutoff. This yielded a definition of persistent waters as sites present at the same location in ≥2 pH values, numbering 1052±197 sites per p*H* (**Fig. 4A**). In contrast, non-persistent (lone) waters, *i.e*., sites found only in one p*H* value showed an upward trend and amounted to 75 at pH 3.5, 177 at pH 4.0, 173 at pH 5.0, 139 at pH 7.0, and 333 at pH 9.0 **(Fig. 4B)**. The most persistent waters (N = 499) were those observed across all five p*H* values (**Fig. 4C**), corresponding to 40.6% of all water molecules observed across all maps. Such a high percentage indicates a localized, well-defined hydration shell that acts as a molecular buffer against p*H*-dependent desolvation and contributes to structural stability, where mostly polar and positively charged residues retain those structural waters (**Fig. 4D**). Lone waters, on the other hand, predominantly appear at higher p*H* (**Fig. 4A**), consistent with total water counts. Analysis of distances between water molecules further revealed that those that persist, cluster substantially closer together than less persistent or lone waters **(Fig. 4E)**. This distribution of distances was maintained across all p*H* conditions **(Fig. S11)**. Our observations imply that persistent water molecules are stabilized by forming water-water hydrogen bonds, in contrast to more isolated, weakly bound waters that are readily exchanged with the bulk solvent. Mapping of these sites on the apoferritin structure solved at p*H* 7 shows a trend where localized negative charge is not likely to organize persistent water molecules **(Fig. 4F)**.

**Figure 4.**
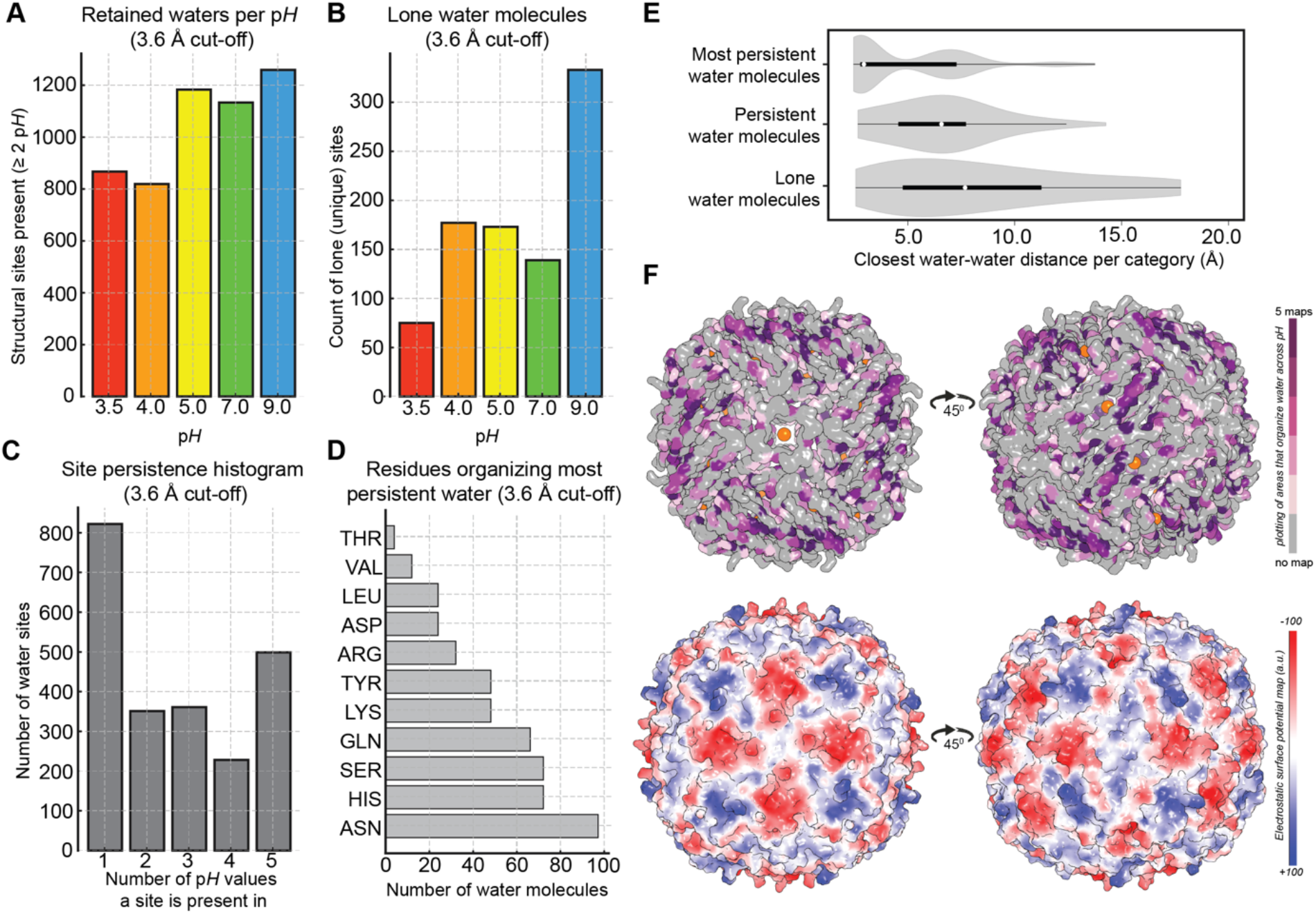
Persistence of water molecules at their sites. (**A**) Retained water molecules per p*H* bin, found in at least one other p*H* map. **(B)** Water molecules per p*H* bin identified in only one map. **(C)** Number of water sites that are identified within a single p*H* map, two, three or four maps, and across all five p*H* maps. **(D)** Water molecules organized across all maps (most persistent) amino acid preferences. Residues not shown do not organize most persistent water molecules. **(E)** Calculated closest water-water distances across the three categories of water molecules, i.e., “lone” water (found only in 1 p*H* map; persistent water, found at least in 2 different p*H* maps; and “most persistent”, found across all resolved maps. **(F)** On top: mapping of persistent sites for the water molecules on top of the biomacromolecule. On the bottom: electrostatic potential map calculated, showing the anti-correlation between persistent water and high electrostatic surface potential.

## Conclusions

Bound water molecules, their nature and adaptability have inspired biochemists, such as Linus Pauling, since the 1950s(*61*). No direct method was available to visualize their physical-chemical adaptations, such as those we report here. Cryo-EM can directly visualize samples of different physical-chemical conditions and interpret stable water molecules that interact with a given biomolecule due to the derived high-resolution. We have quantitatively expanded the 50-year-old hypothesis on protein dehydration as a function of decreasing p*H(4)* and, for the first time, directly visualized and quantified this effect on a biomolecular complex. During this effect, which we term *Kuntz-Kauzmann acid-driven protein desolvation*, the loss of waters leads to a gradual protein destabilization due to missing compensation of protein surface charges by waters (**Fig. 5**).

**Figure 5.**
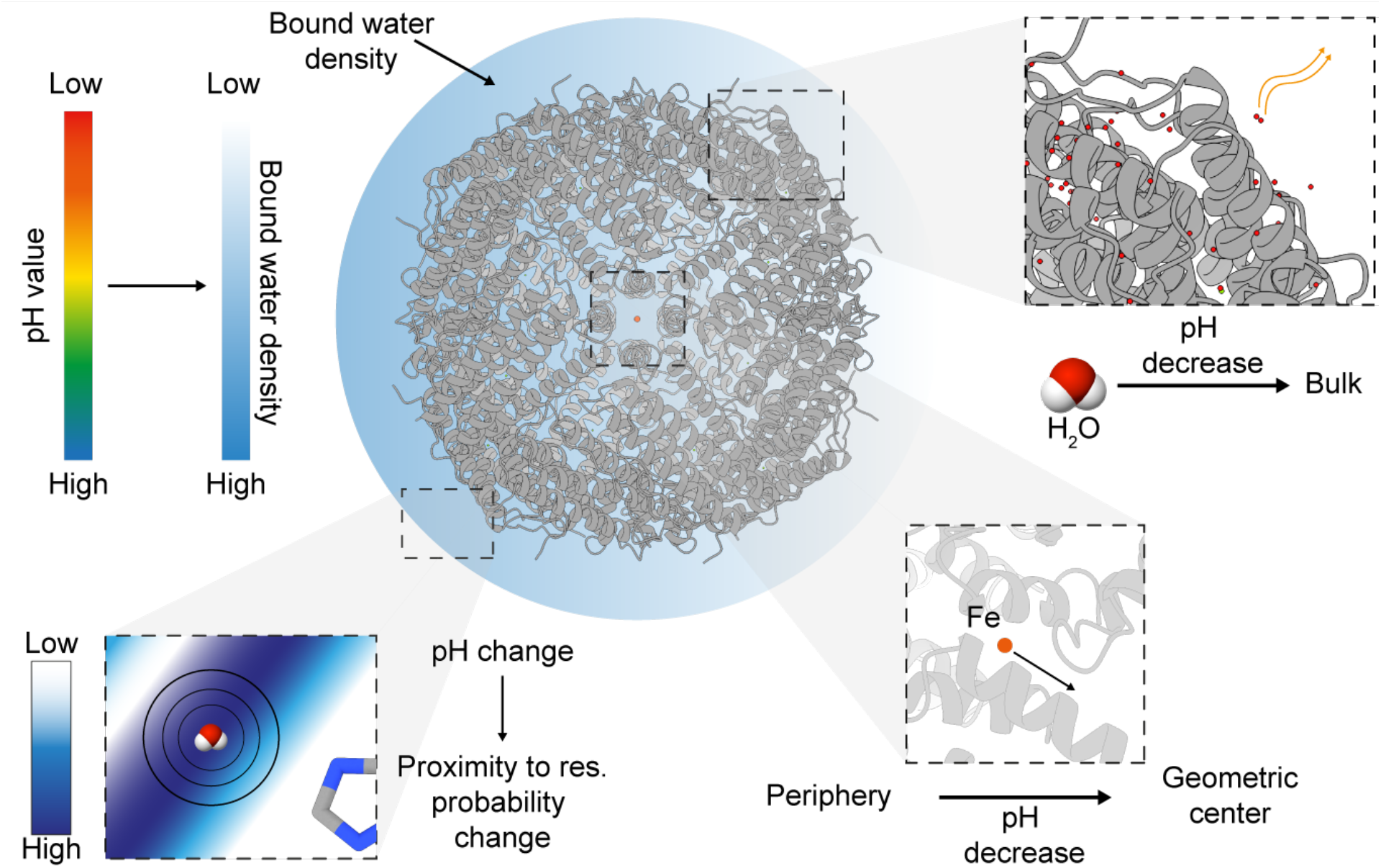
Solvation layer effects as a function of p*H*. The modelled waters belonging to the solvation layer numerically differ with decreasing p*H*. Density of bound waters shows a decrease in accordance with p*H* decrease, moving towards the bulk and not contributing to protein complex stabilization, whereas the probability distribution of water molecule localization in proximity to the nearest amino acid side chain becomes narrower. Fe atoms are affected by water density, being pushed to the geometric center of the protein complex.

This effect is observed across the complete protein surface, and distances of water molecules to the protein surface show broader distributions upon acidification, indicating increased localized instability. Kuntz-Kauzmann acid-driven protein desolvation is more pronounced in proximity to specific side chains, especially negatively charged ones, but also in proximity to lysine but less to arginine and histidine; Lysine is known to change its solvation layer as a function of decreasing p*H*(*58*). However, these residues do not organize stable water molecules; in contrast, the asparagine residues, which are highly solvated across all p*H* values, show the highest degree of proximal structural water molecules that are also retained at their positions. A hypothesis is that their side-chain adaptability, having both partial positive and negative charges, may substantially enhance the localized attraction of stable water molecules.

Overall, we find that the p*H*-dependence of resolved structural waters is described by two p*H*-responsive site classes that titrate around acidic and basic conditions, and a third p*H*-independent “persistent shell”. The three classes constitute a coarse-grained envelope of proton-linked binding in Wyman’s framework(*62*) where negative linkage explains loss of waters upon acidification. Additionally, our MD shows that deprotonation at acidic sites lengthens water residence times from a few picoseconds to ∼10 ps, corresponding to ∼0.5 kcal·mol^−1^ per site. When considered across total hydration sites, values are compatible with local charge-regulation energetics(*63*).

Conclusively, our observations and experimental set-up utilizing broad range buffer series for the study of biological macromolecules in variable p*H*, combined with molecular dynamics simulations to get insight into residence times, accelerates understanding of solvation layer effects on biological macromolecules. Our results can drive applications in lead compound development for active sites (e.g., the observation of iron displacement with decreasing p*H*) or allosteric sites where water molecules play an active structural/functional role. Finally, there is an increasing demand for biotechnological protein applications at low p*H* (e.g., extraction of metals or rare earth elements(*64*)). Our statistical observations based on the high-resolution structures of a protein complex can also act as a lead for protein design of highly soluble and stable proteins, informing the current era of machine-learning-based protein design(*65*).

## Methods

### Protein expression and purification

The genomic sequence for human apoferritin was present in a pGEX2T vector, containing a N-terminal GST-fusion and modified to contain a TEV cleavage site, instead of a thrombin cleavage site, resulting in Plasmid LF2422. This plasmid was kindly provided by Dr. Louise Fairall and Dr. Christos Savva (Protex facility at the University of Leicester). Protein expression and purification was achieved, following a modified protocol from(*66*). Briefly overexpression of human apoferritin was achieved using Escherichia coli BL21 Rosetta™ 2(DE3) (Novagen) transformed with plasmid LF2422. Cells were grown at 37 °C until A600 = 1.0 before adding 1 mM 1-thio-b-d-galactopyranoside (IPTG) and shifting the temperature to 20 °C. After growth overnight, cells were harvested by centrifugation (2967 x g) at 4 °C, for 30 min. The pellet was resuspended in 100 mL / L (bacterial culture) of phosphate buffered saline (PBS), containing 0.5 % Triton X-100 and 0.5 mM tris(2-carboxyethyl) phosphine (TCEP) and Roche Complete protease-inhibitors. Cells were lysed using a continuous cell disruptor (Constant Systems), and the lysate was centrifuged at > 100000 x *g*, 4 °C for one hour.

Cleared lysate was initially purified using a 5 mL GSTrap HP column (Cytiva). After binding and washing the column with 5 CV of washing buffer (1 x PBS, 0.5 % Triton X-100, 0.5 % TCEP) a buffer exchange to the SEC buffer (5 CV, 50 mM Tris, pH 7.5, 100 mM NaCl, 0.5 mM TCEP) and on-column digestion of the bound protein was performed with 100 µg self-prepared TEV-protease. After flushing the cleaved protein from the column, the bound GST-tag and any non-cleaved protein were eluted from the column using the elution buffer (50 mM Tris, pH 7.5, 100 mM NaCl, 0.5 mM TCEP, 10 mM reduced Glutathione). Cleaved apoferritin was then further purified using a Phenomenex BioSep SEC-s4000. After size-exclusion chromatography, the peak with fractions containing fully assembled apoferritin (MW: 450 kDa) was pooled.

### pH Change, cryo-preparation, vitrification and data acquisition

The pool was concentrated to about 500 µL using a Millipore 100 kDa cutoff Amicon tube. The buffer-exchange and further concentration was performed using Millipore 100 kDa cutoff tube filters. Buffers for p*H*-change were prepared, following the procedure described elsewhere(*46*). In brief, buffers were prepared by mixing two stock solutions in different volumes. Stock solution A contained 200 mM boric acid and 50 mM citric acid. Stock solution B contained 100 mM Na_3_PO_4_. Surplus protein sample was frozen in liquid nitrogen and stored at −80 °C. All different p*H* samples (3.5, 4, 5, 7, 9) were diluted to 0.2 mg/ml prior to vitrification.

For cryo-EM sample preparation, R2.1 200-mesh holey carbon grids (Quantifoil Micro Tools) were glow-discharged using a PELCO easiGlow system (15 mA, 25 s, mbar residual air pressure). Approximately 3.5 μL of each sample was applied to the grids in a Vitrobot Mark IV (Thermo Fisher Scientific) operated at 100% relative humidity and 4 °C. Grids were blotted for 4 s using 595-grade ashless filter paper (Thermo Fisher Scientific) with blot force 0 and immediately plunge-frozen into liquid ethane. Vitrified grids were transferred under cryogenic conditions to our 200 keV Thermo Fisher Scientific Glacios transmission electron microscope equipped with a Falcon IIIEC direct electron detector. For each of the five p*H* conditions, ∼2,000–4,000 movies were recorded (see **Table S1**) at a calibrated pixel size of 0.59 Å/pixel, with a total accumulated electron dose of 30 e−/Å^2^ fractionated across 30 frames.

### Image processing of cryo-EM data, model building and refinement

All image processing steps were performed within the framework of SCIPION v3.1.0(*67*). In detail, each of the collected datasets (pH 3.5: 3,725 images, pH 4: 2,308 images, pH 5: 2,541 images, pH 7: 1,891 images, pH 9: 1,867 images) was imported, corrected for beam-induced motion with RELION motioncorr(*68*), and CTF parameters were calculated with gctf(*69*). Manual particle picking of ∼1,000 particles followed, and this particle set was used for TOPAZ training and picking(*70*). In parallel, picking was also performed with xmipp3(*71*), as well as crYOLO(*72*) and a consensus pick was derived from all different picking methods. After combining the multiple picking strategies, each dataset yielded an initial particle set of (pH 3.5: 710,073 particles, pH 4: 280,270 particles, pH 5: 521,922 particles, pH 7: 529,107 particles, pH 9: 438,640 particles) which was subsequently refined by iterative 2D classification with 300 classes in cryoSPARC v3.2, leading to final particle sets of (pH 3.5: 217,187 particles, pH 4: 267,457 particles, pH 5: 421,029 particles, pH 7: 432,247 particles, pH 9: 343,262 particles). The refined particle sets were employed in RELION-4(*73*) 3D initial modeling, followed by cryoSPARC 3D non-uniform refinement of the *ab-initio* models, taking into consideration per-particle motion and CTF parameters, and then the final particle sets and volumes were imported again to RELION-4(*73*) for Bayesian polishing, leading to final reconstructions of 1.99 Å, 2.06 Å, 1.96 Å, 2.08 Å and 1.89 Å (Gold-standard FSC = 0.143) for pH 3.5, 4, 5, 7, and 9 respectively.

The amino-acid sequence of the *H. sapiens* ApoF heavy chain (Uniprot id: P02794) monomer was fit into each of the different pH EM reconstructions using ChimeraX(*74*), and then real space refined in PHENIX(*29*). After automatic refinement, each ApoF monomer model was manually inspected and side-chain fits were refined in COOT v0.9.2(*75*). Modelling of waters was performed in two steps. First, each pH map and its corresponding model were inspected in coot and waters were manually modeled, taking into consideration physical-chemical restraints. Simultaneously, each different pH model-map combination was used as input in the DOUSE module of PHENIX, for automated water placement in bulk, with cutoffs from 2 to 4 A. Subsequently, the monomer density with the added waters was once again manually inspected in COOT, with a uniform sigma level (0.5), and water placement was refined manually as needed, comparing automated and manual water placement in the models. For the generation of the complete ApoF complex, the find_symmetry and apply_NCS_symmetry modules in PHENIX were used. Finally, all duplicated water molecules were eliminated at the symmetry axis points of the model by employing a custom Python script.

### Constant pH (CpH) MD simulations

The cryo-EM structures of apoferritin, obtained at p*H* 4, 7 and 9, were used as starting models. For each structure, seven neighboring protomers were extracted to build a minimal system for running CpHMD simulations (**Fig. S12**). The phbuilder tool(*76*) was used to prepare the CpHMD simulations for the GROMACS software. Titration was allowed only for the glutamate, aspartate, histidine and lysine residues of the middle monomer as well as for residues of neighboring monomers that can directly interact with the middle monomer. Residues coordinating the iron(II) and magnesium(II) ions were kept at their deprotonated state (for E24, E59) or neutral state (for HID62, HIE170) in order to retain the binding site of these ions. Further information about the titrated residues for each monomer can be found in the supporting information (**Table S3**). The system was solvated in a box with a border of a minimum distance of 1 nm to the protein and neutralized with NaCl and buffer molecules. The topology was generated using the CHARMM36-mar2019-cphmd force field(*77*) in combination with the CHARMMff additive force field for iron(II)(*78*) and TIP3P water model. After minimization using the steepest descent algorithm, the system was further equilibrated in an NVT ensemble and subsequently in an NPT ensemble, while gradually releasing restraints on the heavy atoms. Both equilibration steps were run for 10 ps using a time step of 1 fs with the temperature regulated by the V-rescale thermostat at 310 K and the pressure kept at 1 bar using the C-rescale barostat, respectively. The PME method was used to perform electrostatics, and the LINCS-algorithm was used to constrain bond lengths. Titration MD runs were performed for 40 ns with a timestep of 2 fs using the GROMACS CpHMD beta software, obtained from https://gitlab.com/gromacs-constantph/constantph. A position restraint of 50 kJ mol^-1^ nm^-2^ was applied to all backbone heavy atoms of the six outer monomers and the iron(II) ion in all simulations. The starting titration states were chosen according to the standard pKa values of the residues and the titration was performed in steps of 0.5 from pH 1.0 to pH 10.5. Titration curves were fitted using the Scipy module(*79*) after calculating the protonation fractions according to equation (1) with *n_deprot_* including all lambda values smaller than 0.2 and *n_prot_* higher than 0.8 using the accompanying script from the GROMACS constant-p*H* Gitlab repository (https://gitlab.com/gromacs-constantph).

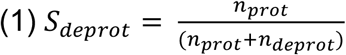

### Whole-assembly apoferritin MD simulations

For all determined cryo-EM structures (pH 3.5, pH 4, pH 5, pH 7, pH 9), MD simulations of the whole 24-mer assembly were run in triplicate with the protonation states of titratable residues adjusted according to the results of the CpHMD simulation. This time, titration states were not allowed to change within one simulation. The protein structures were placed in a cubic box with 1 nm distance between protein and box and subsequently solvated with TIP3P water. NaCl was used to neutralize the system and to ensure an ionic strength of 150 mM. The minimization and equilibration steps were performed as described above, for 100 ps in both NVT and NPT ensembles. Production runs were conducted for 21 ns in triplicates each using GROMACS version 2021 (https://doi.org/10.5281/zenodo.4457626).

### Hydration site and residence time analysis

Following 20 ns of water equilibration around the full apoferritin complex, 500 evenly spaced frames were taken from the last ns of each simulation and used for trajectory analysis. Rotational and translational movements of the protein were removed by aligning the whole trajectory to the apoferritin structure in the first frame. A water density map of the regions up to 5 Å around the protein with a grid spacing of Å was calculated using the MDAnalysis package(*80*). This map was smoothed by calculating the mean water density of each grid cell and its adjacent grid cells that were sharing an edge. Close density peaks with a distance smaller than 1 Å between them were averaged using the DBscan algorithm(*81*) implemented in the Sklearn package(*82*). Hydration sites were counted if they had a smoothed density of 6 kg/L or more. Closest residues were defined as the residues that most often had the closest heavy atom distance to the hydration site. The water residence times for all glutamates were calculated by integrating the survival probability function of water 3.6 Å around each respective glutamate residue using MDAnalysis(*80*).

## Supporting information

Supplementary material including 12 Figures and 3 Tables

## Acknowledgements

The authors would like to deeply thank Dr. Christos Savva (Diamond Light Source) that provided the plasmid and Prof. Milton T. Stubbs (University of Halle) for initiating the scientific discussions on the correlation of p*H* dependence and charge distribution of cryo-EM maps. P.L.K. was supported by the European Union through funding of the Horizon Europe ERA Chair “hot4cryo” project no. 101086665, the Federal Ministry of Education and Research (BMBF, ZIK program) (grant nos. 03Z22HN23, 03Z22HI2, and 03COV04), the European Regional Development Funds (EFRE) for Saxony-Anhalt (grant no. ZS/2016/04/78115 and ZS/2024/05/187255), the Deutsche Forschungsgemeinschaft (project nos. 391498659, RTG 2467 and 514901783, SFB 1664 [A04, C04, and D01]), and the Martin Luther University Halle-Wittenberg. G.K. acknowledges funding from DFG (project no. 514901783, SFB 1664, subproject D01). The authors further thank the high-performance computing centers at Leipzig University and at the NHR center of TU Dresden for providing the computational resources. D.A.S. was supported by FCT - Fundação para a Ciência e a Tecnologia, I.P., through MOSTMICRO-ITQB. R&D Unit (DOI 10.54499/UIDB/04612/2020; DOI 10.54499/UIDP/04612/2020;) and LS4FUTURE Associated Laboratory (DOI 10.54499/LA/P/0087/2020). I.S. acknowledges funding from the European Union HORIZON-MSCA-2024-PF-01 under grant agreement no. 101207412 (Protein ER-ways).

## Contributions

J.M. performed sample preparation and biochemical assays together with F.L.K. F.L.K. prepared cryo-EM grids and carried out screening together with F.H. F.H. collected the high-resolution cryo-EM data. J.B. conceived, developed, and implemented the initial pipeline for water analysis. D.A.S. processed the cryo-EM micrographs, deriving the maps. I.S. performed structural modelling with support from C.T. P.L.K. carried out data analysis and distance calculations. P.L.K., F.H., and I.S. interpreted the data and evaluated the results. I.K.S. performed molecular dynamics simulations under the supervision of G.K., and together they evaluated the outcomes. P.L.K. wrote the manuscript with input from all authors. P.L.K. and F.H. conceived the project, with guidance from G.K. regarding the molecular dynamics. P.L.K. supervised the project, and both P.L.K. and G.K. acquired funding.

## Author Contributions (CRediT taxonomy)

Conceptualization: P.L.K., F.H., G.K. Methodology: J.M., F.L.K., J.B., D.A.S., I.S., C.T., I.K.S. Software: J.B., D.A.S. Validation: P.L.K., F.H., I.S., I.K.S., G.K. Formal analysis: P.L.K., I.S., I.K.S. Investigation: J.M., F.L.K., F.H., D.A.S., I.S., C.T., I.K.S. Resources: P.L.K., G.K. Data curation: D.A.S., P.L.K., I.S. Writing - Original Draft: P.L.K. Writing - Review & Editing: All authors. Visualization: P.L.K., I.S., F.H. Supervision: P.L.K., G.K. Project administration: P.L.K. Funding acquisition: P.L.K., G.K.

## Competing interests

The authors declare no competing interests.

## Data availability statement

Cryo-EM maps and models were deposited in EMDB and PDB, respectively, and correspond to the following entries: p*H* 3.5: EMD-54951, 9SJR; p*H* 4.0: EMD-54952, 9SJS; p*H* 5.0: EMD-54953, 9SJT; p*H* 7.0: EMD-54954, 9SJU; p*H* 9.0: EMD-54955, 9SJV.

## Notes

### Competing Interest Statement

The authors have declared no competing interest.

