## Supplementary material including 12 Figures and 3 Tables for "Direct evidence of acid-driven protein desolvation"

### Supplementary Figures

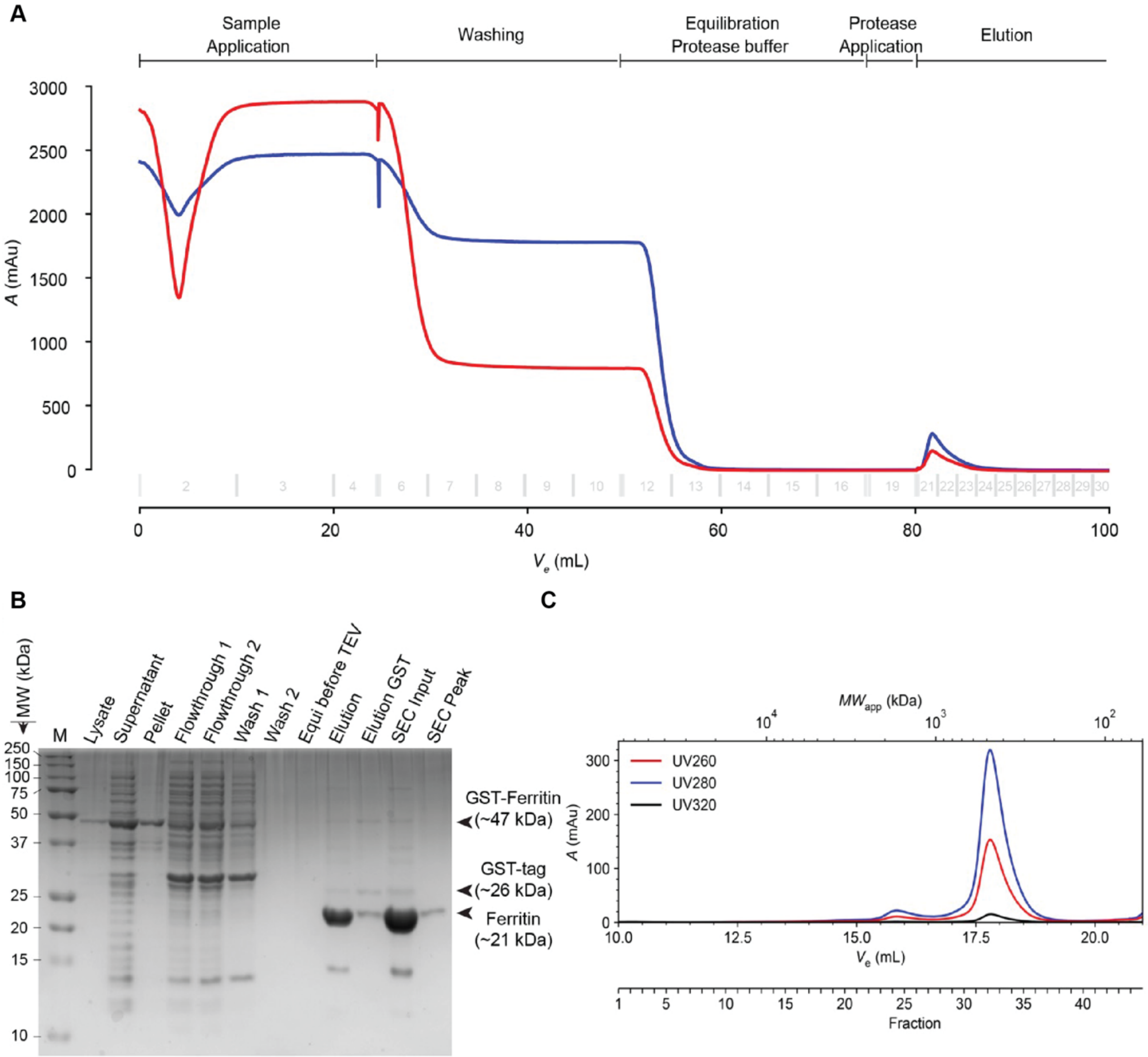


**Figure S1.** Expression and Purification. (**A**) Chromatogram of the purification of apoferritin from bacterial lysate. Purification phases are noted above the chromatogram and UV-absorption at 260 nm and 280 nm are drawn in red and blue respectively. The first peak after on-column digestion with TEV-protease was further purified using size-exclusion chromatography. (**B**) SDS-PAGE of the complete purification of apoferritin. GST-Ferritin, the GST-tag after TEV-cleavage and cleaved apoferritin are marked with arrows at their respective MW. **C:** SEC-Profile of cleaved apoferritin. UV-absorption at 260 nm, 280 nm and 320 nm are drawn in red, blue and black respectively and the apparent molecular weight is shown in a logarithmic scale above the chromatogram. The peak containing fully assembled apoferritin (Fractions 32 to 36, MW 400-500 kDa) was collected and subjected to pH and vitrification.

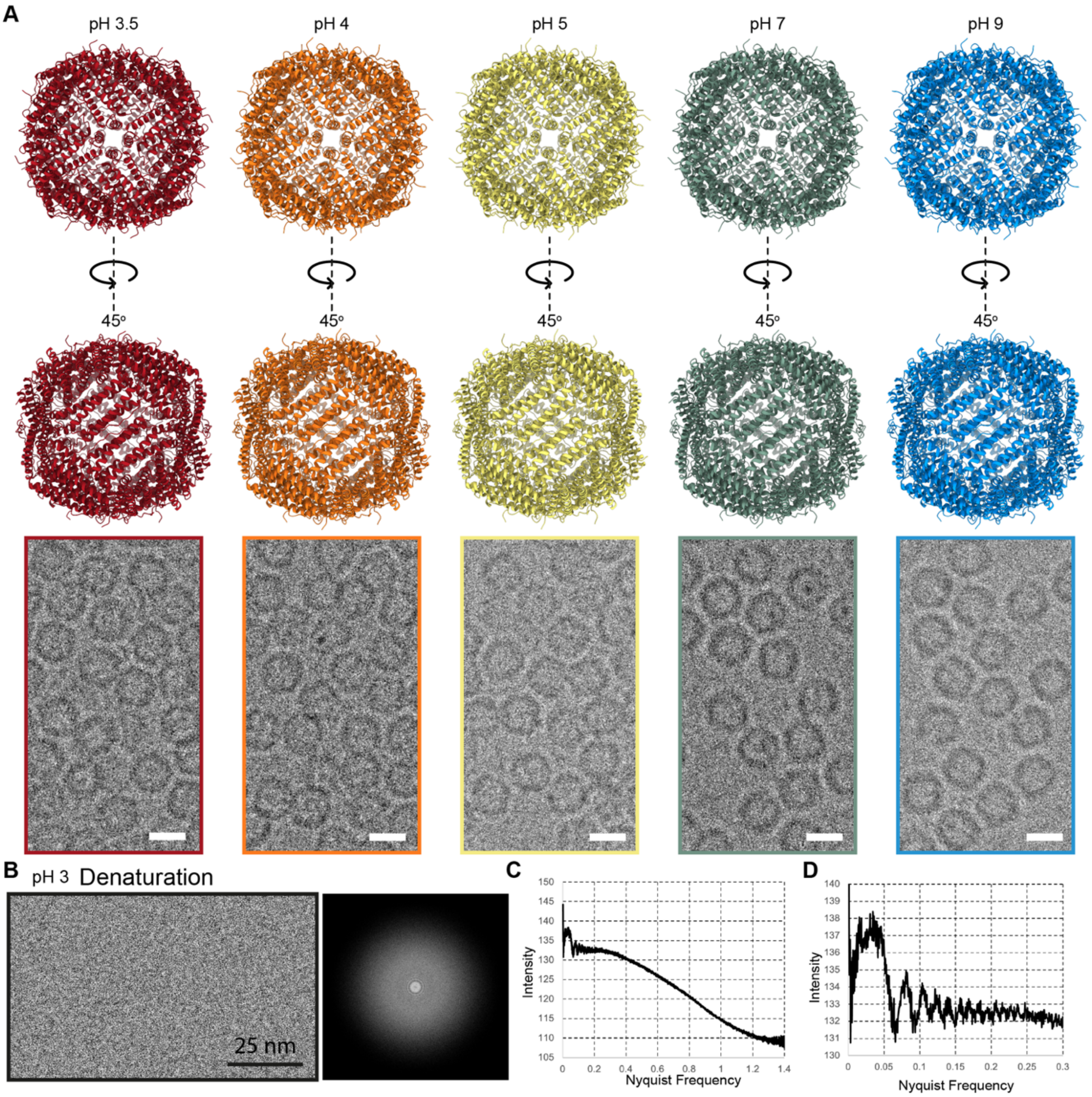


**Figure S2.** Apoferritin models and representative micrographs across p*H* values. (**A**) Resulting model and data collection representative micrograph for ApoF cryo-EM data collection at pH 3.5., pH 4, pH 5, pH 7 and pH 9. (B) At pH =< 3 there are no observable intact apoferritin particles visible on the cryo-EM micrographs, but the Fourier transform of the image shows protein presence. (C-D) Quantitative analysis of Fourier Transform (FT) intensities show oscillations at different Nyquist frequencies, indicating presence of proteinaceous material. Scale bars = 10 nm.

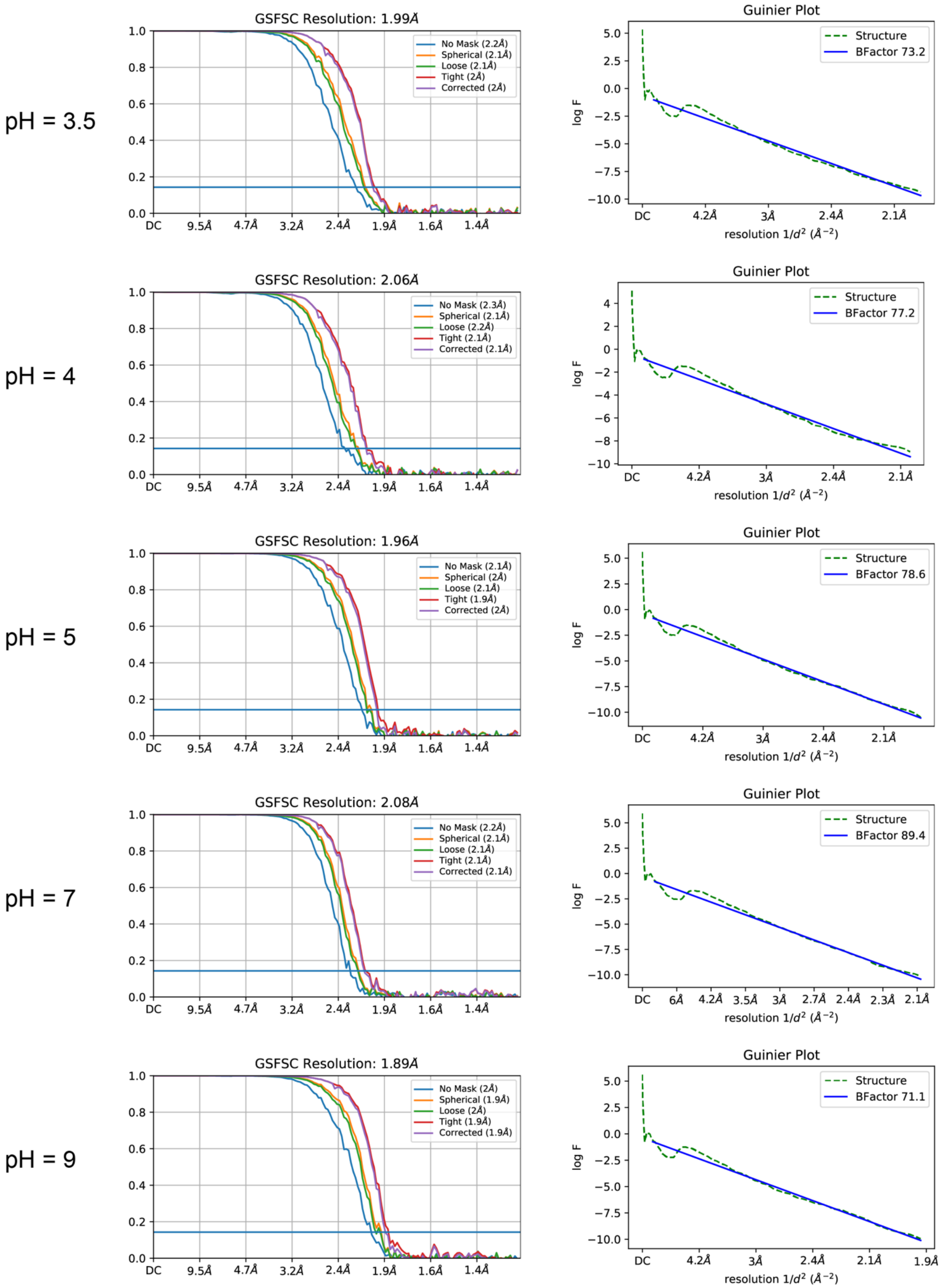


**Figure S3**. Fourier Shell Correlation plots for each reconstruction**.** FSC (Fourier Shell Correlation) and Guinier Plots for all reconstructions at the different p*H*s.

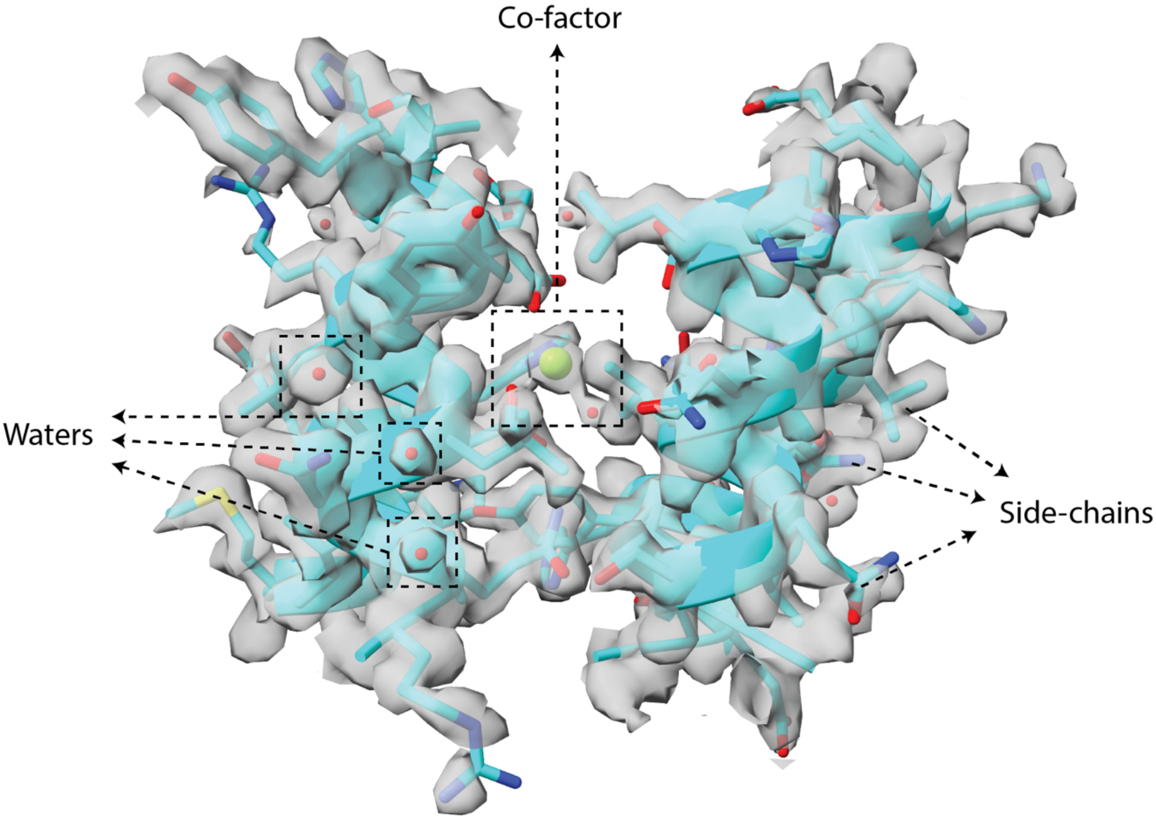


**Figure S4.** Resolvability. Example of resolvability for different molecule classifications in resolved cryo-EM maps. In all resolved structures, water molecules, co-factors (*e.g.,* magnesium) and side-chain densities were clearly visible and able to be modeled accordingly. Map threshold is 1.07, step 1.

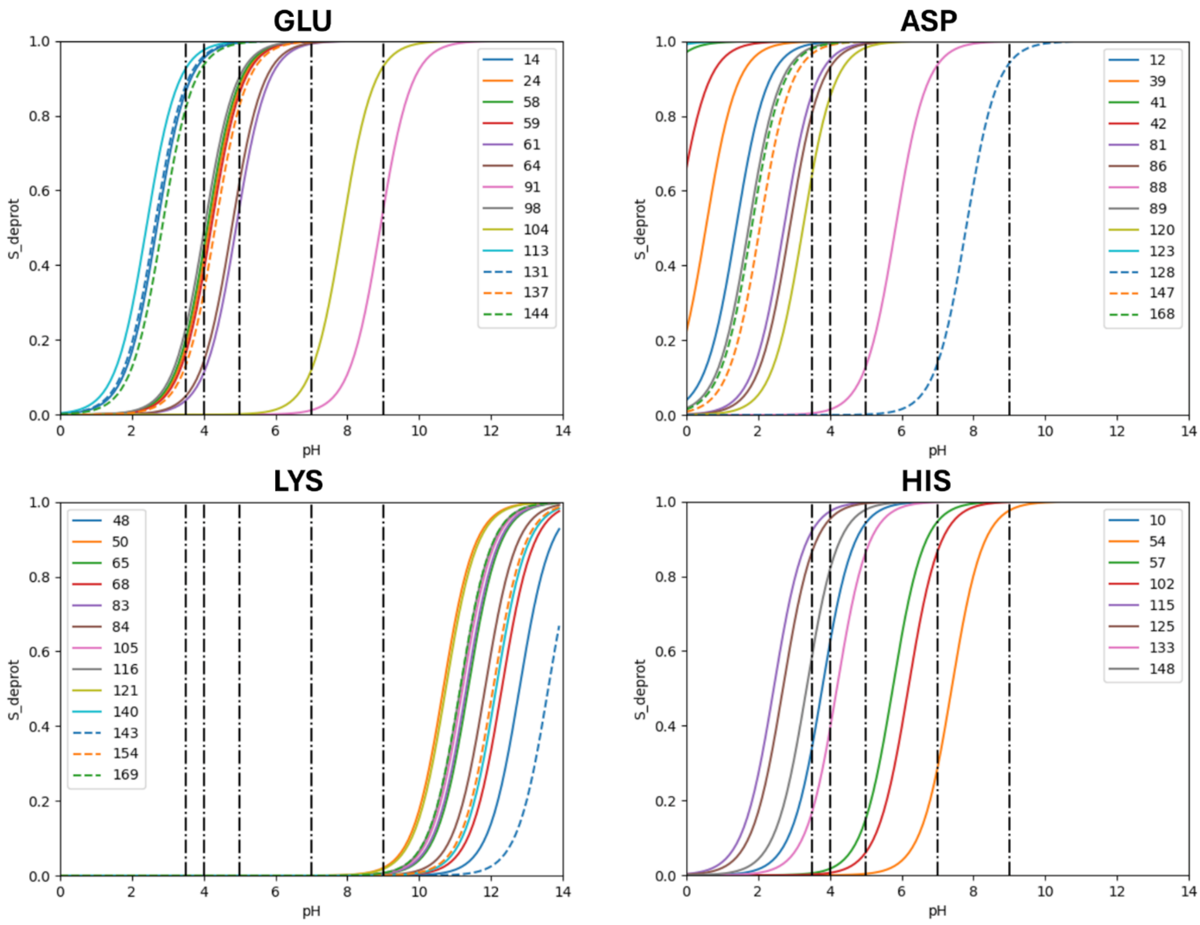


**Figure S5.** Calculated titration curves for the side-chain functional groups of Asp, Glu, Lys and His residues in the apoferritin structure calculated by CpH MD simulations. Vertical lines indicate the different pH values which should be adjusted in the whole-assembly apoferritin MD simulations.

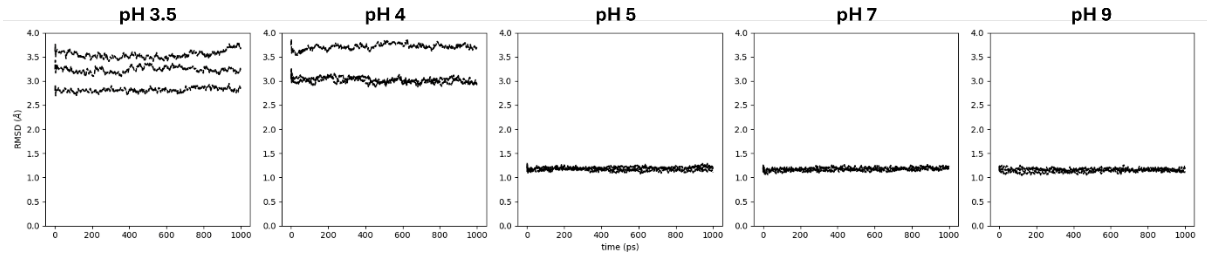


**Figure S6.** RMSD-vs-time plots of the whole apoferritin complex relative to the cryo-EM structure in the MD simulations at different p*H* values**.**

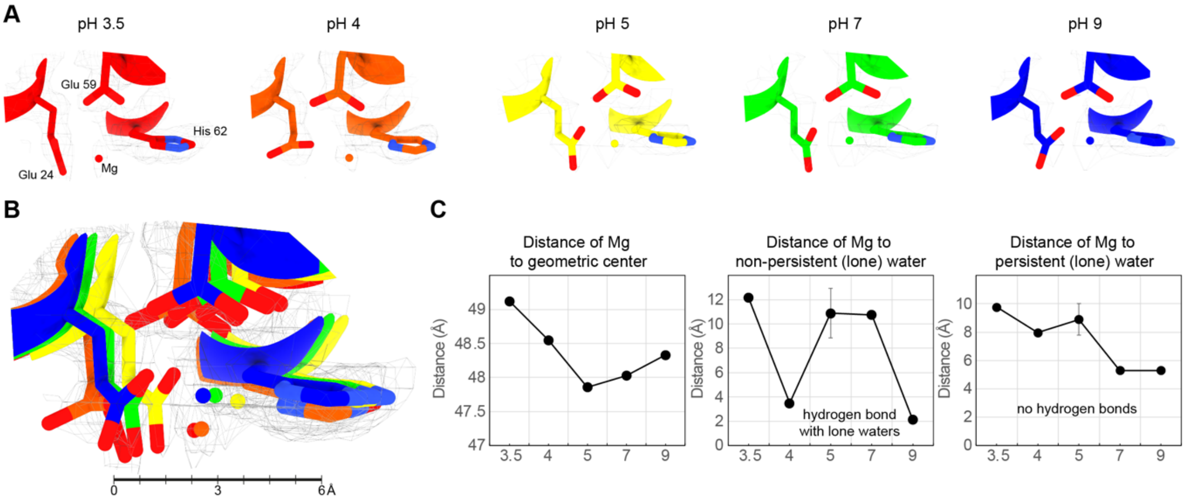


**Figure S7.** Contacts of the magnesium ion. **(A)** The magnesium ion remains stable as function of p*H*. **(B)** Local superposition of residues organizing the magnesium, and the 3D location of the ion, color-coded as function of pH. Its movement is minimal. **(C)** he magnesium shows a complex movement in respect to the geometric center (left plot), and hydrogen bonds only occur with "lone" water molecules, i.e., water molecules recovered in only a single p*H*. Right plot shows that Mg does not form hydrogen bonds with persistent water molecules as opposed to iron.

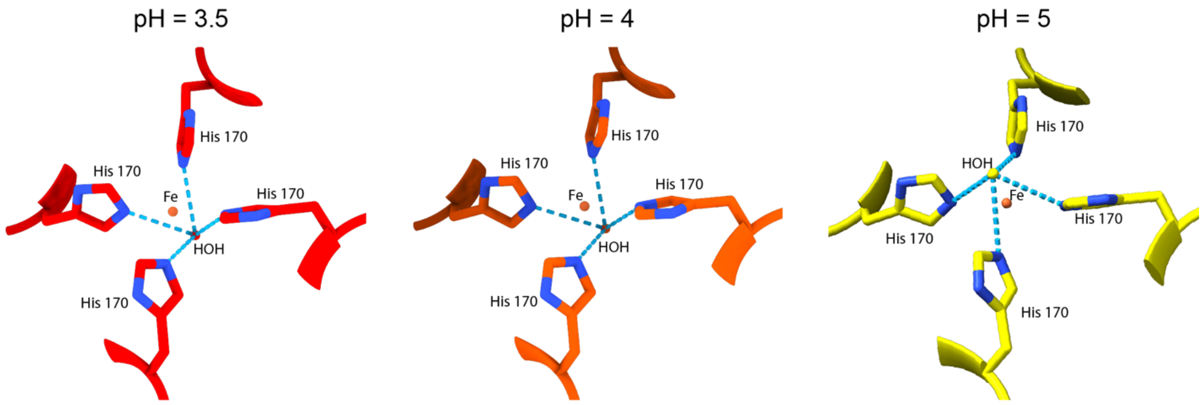


**Figure S8.** Interactions of the water molecule in proximity to the iron with surrounding histidines.

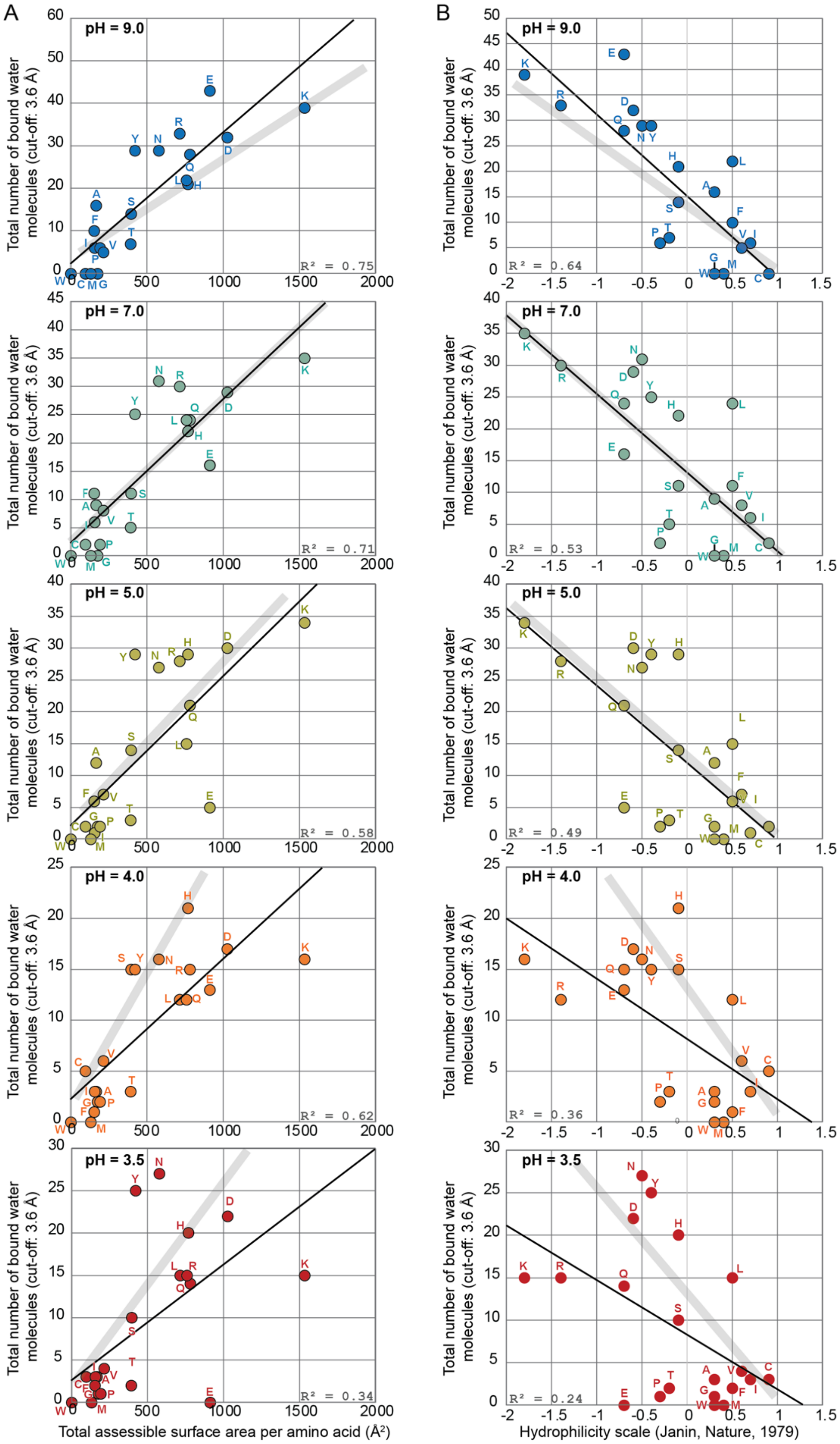


**Figure S9.** Accessible surface area are strong predictors of protein solvation, but such descriptors are increasingly weaker with lower p*H*. Grey line represents the trend observed at p*H* 7. Note that values in x and y axes are aggregate numbers per amino acid.

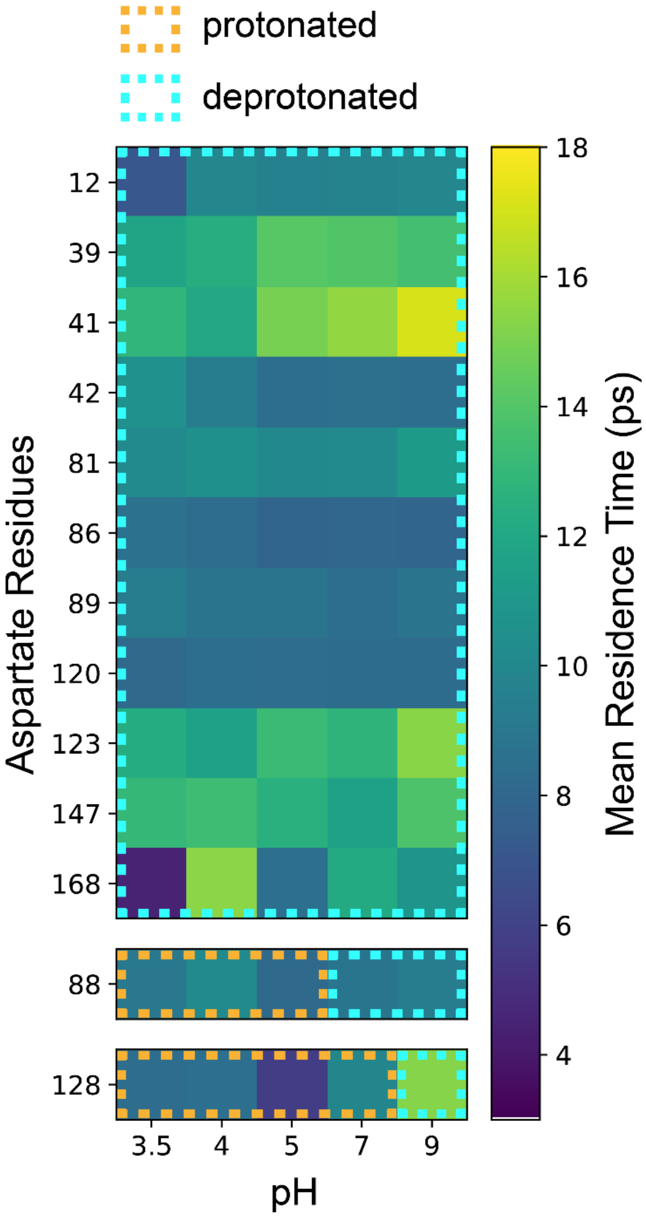


**Figure S10.** Residence times of waters surrounding the aspartate residues at different p*H* values.

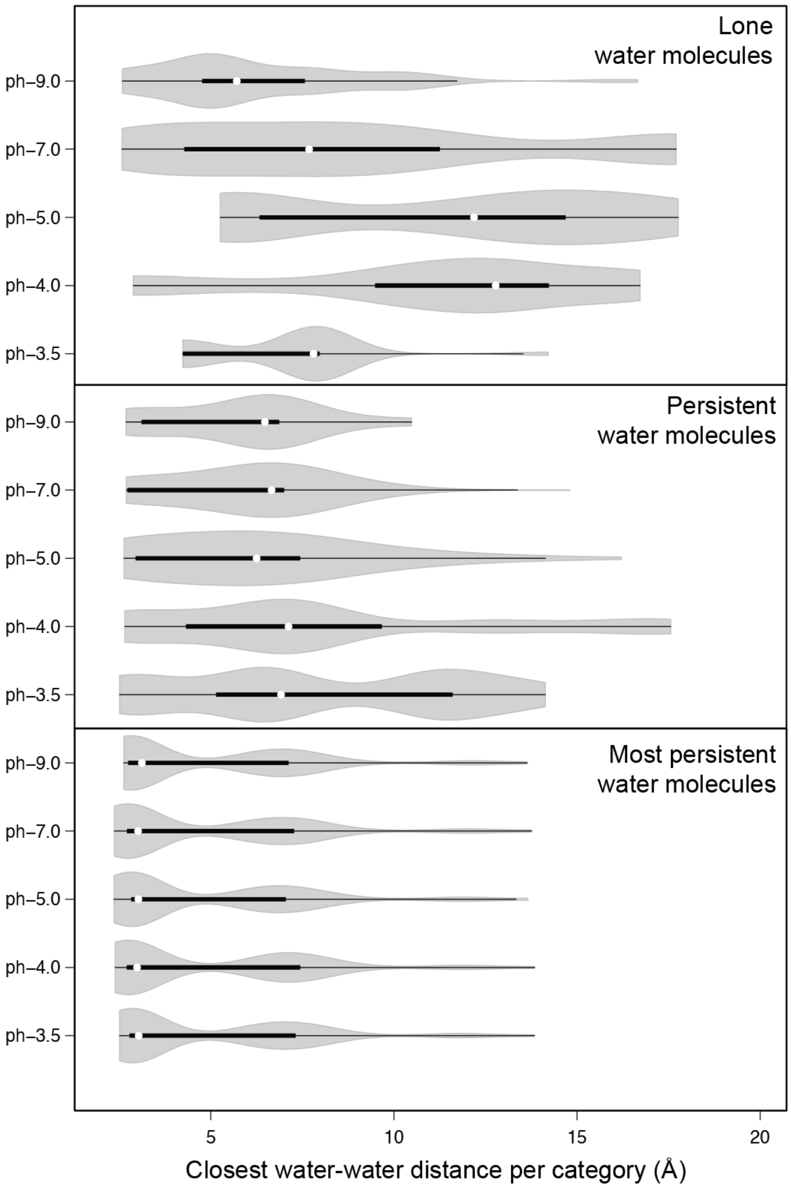


**Figure S11.** Distribution of closest water-water distances for different categories of water molecules across p*H* conditions. Violin plots show the distribution of minimum distances (Å) between water molecules classified as (top) lone water molecules, (middle) persistent water molecules, and (bottom) most persistent water molecules at pH 3.5, 4.0, 5.0, 7.0, and 9.0. White dots represent the median values, thick black bars indicate the interquartile range, and thin black lines show the 95% confidence intervals. The width of the gray violin plots reflects the density of observations at each distance value.

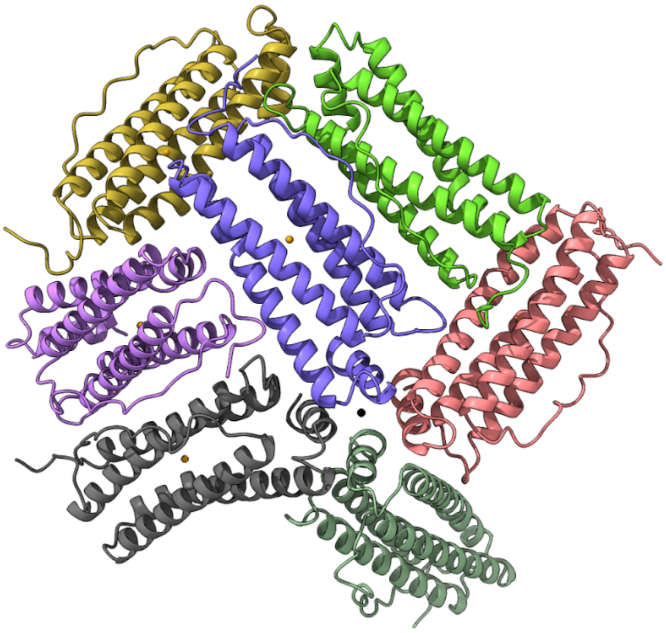

**Figure S12.** Apoferritin 7-mer system for running CpH MD simulations. The seven protein chains are displayed with different colors. A – dark purple, B - fuchsia, C - dark green, H - black, N - purple, P - yellow, Q – bright green. Magnesium and iron ions are colored orange and black, respectively. The titrated residues are listed in **Table S2**.

### Supplementary Tables

**Table S1**. Cryo-EM data collection, refinement and validation statistics.

| | #1 p*H* 3.5 (EMDB-54951) (PDB 9SJR) | #2 p*H* 4 (EMDB-54952) (PDB 9SJS) | #3 p*H* 5 (EMDB-54953) (PDB 9SJT) | #4 p*H* 7 (EMDB-54954) (PDB 9SJU) | #5 p*H* 9 (EMDB-54955) (PDB 9SJV) |
|---|---|---|---|---|---|
| **Data collection and processing** | | | | | |
| Magnification | 240,000 | 240,000 | 240,000 | 240,000 | 240,000 |
| Voltage (kV) | 200 | 200 | 200 | 200 | 200 |
| Image Nr. | 3,725 | 2,308 | 2,541 | 1,891 | 1,867 |
| Electron exposure (e–/Å$^2$) | 30 | 30 | 30 | 30 | 30 |
| Defocus range (μm) | 1.0-2.0 μm | 1.0-2.0 μm | 1.0-2.0 μm | 1.0-2.0 μm | 1.0-2.0 μm |
| Pixel size (Å) | 0.59 | 0.59 | 0.59 | 0.59 | 0.59 |
| Symmetry imposed | O | O | O | O | O |
| Initial particle images (no.) | 710,073 | 280,270 | 521,922 | 529,107 | 438,640 |
| Final particle images (no.) | 217,187 | 267,457 | 421,029 | 432,247 | 343,262 |
| Map resolution (Å) FSC threshold | 1.99 (0.143) | 2.06 (0.143) | 1.96 (0.143) | 2.08 (0.143) | 1.89 (0.143) |
| **Refinement** | | | | | |
| Map sharpening *B* factor (Å$^2$) | 73.2 | 77.2 | 78.6 | 89.4 | 71.1 |
| Model composition | | | | | |
| Non-hydrogen atoms | 35016 | 35088 | 35304 | 35352 | 35664 |
| Protein residues | 4152 | 4152 | 4152 | 4152 | 4152 |
| Ligands | 48 | 48 | 48 | 48 | 48 |
| Waters | 960 | 1032 | 1248 | 1296 | 1608 |
| *B* factors (Å$^2$) | | | | | |
| Protein | 4.93/41.53/14.10 | 7.59/40.25/17.21 | 2.74/33.02/10.01 | 5.29/41.13/14.68 | 1.01/28.35/5.02 |
| Ligand | 5.08/24.64/14.86 | 8.38/27.77/18.07 | 8.06/13.43/10.75 | 12.17/20.10/16.14 | 2.08/4.77/3.42 |
| Water | 5.80/30.00/15.12 | 8.98/25.00/17.73 | 6.31/19.35/10.88 | 9.30/30.00/15.99 | 1.57/20.00/6.26 |
| R.m.s. deviations | | | | | |
| Bond lengths (Å) | 0.004 | 0.004 | 0.004 | 0.004 | 0.004 |
| Bond angles (°) | 0.904 | 0.886 | 0.889 | 0.938 | 0.898 |
| Validation | | | | | |
| MolProbity score | 1.36 | 1.45 | 1.70 | 1.55 | 1.32 |
| Clashscore | 5.12 | 8.26 | 7.37 | 7.03 | 3.67 |
| Poor rotamers (%) | 1.30 | 0.65 | 1.95 | 1.30 | 1.30 |
| Ramachandran plot | | | | | |
| Favored (%) | 98.83 | 98.25 | 97.66 | 97.66 | 97.66 |
| Allowed (%) | 1.17 | 1.75 | 2.34 | 2.34 | 2.34 |
| Disallowed (%) | 0.00 | 0.00 | 0.00 | 0.00 | 0.00 |

**Table S2**. Kolmogorov-Smirnov (KS) statistics and summary of water–protein distance distributions. Pairwise comparisons were performed between always present, persistent (sometimes present), and lone (single-site) water molecules across all p*H* values and within each p*H* condition (3.5, 4.0, 5.0, 7.0, 9.0). For each comparison, the KS statistic and *p-value* are shown, as well as mean and median distances for both groups ($Mean_1/Mean_2$, $Median_1/Median_2$), sample sizes ($N_1/N_2$), and standard deviations ($SD_1/SD_2$). Global values (p*H* all) correspond to the aggregate distributions of all waters across all conditions.

| pH | Comparison | KS statistic | *p-value* | $Mean_1$ | $Mean_2$ | $Med_1$ | $Med_2$ | $N_1$ | $N_2$ | $SD_1$ | $SD_2$ |
|---|---|---|---|---|---|---|---|---|---|---|---|
| all | always vs persistent | 0.387 | $5.0\times10^{-44}$ | 5.00 | 6.44 | 2.95 | 6.58 | 499 | 940 | 2.80 | 2.51 |
| all | always vs lone | 0.439 | $1.7\times10^{-54}$ | 5.00 | 8.48 | 2.95 | 7.68 | 499 | 822 | 2.80 | 4.45 |
| all | persistent vs lone | 0.294 | $8.1\times10^{-34}$ | 6.44 | 8.48 | 6.58 | 7.68 | 940 | 822 | 2.51 | 4.45 |
| 3.5 | always vs persistent | 0.365 | $1.2\times10^{-25}$ | 5.05 | 7.97 | 3.03 | 6.92 | 499 | 368 | 2.78 | 3.78 |
| 3.5 | always vs lone | 0.561 | $1.1\times10^{-19}$ | 5.05 | 6.97 | 3.03 | 7.80 | 499 | 75 | 2.78 | 2.25 |
| 3.5 | persistent vs lone | 0.389 | $5.5\times10^{-9}$ | 7.97 | 6.97 | 6.92 | 7.80 | 368 | 75 | 3.78 | 2.25 |
| 4.0 | always vs persistent | 0.354 | $3.0\times10^{-22}$ | 5.05 | 8.00 | 2.98 | 7.12 | 499 | 320 | 2.86 | 4.62 |
| 4.0 | always vs lone | 0.732 | $2.6\times10^{-69}$ | 5.05 | 11.14 | 2.98 | 12.78 | 499 | 177 | 2.86 | 4.31 |
| 4.0 | persistent vs lone | 0.534 | $2.3\times10^{-30}$ | 8.00 | 11.14 | 7.12 | 12.78 | 320 | 177 | 4.62 | 4.31 |
| 5.0 | always vs persistent | 0.245 | $4.6\times10^{-15}$ | 4.98 | 6.04 | 3.02 | 6.25 | 499 | 615 | 2.76 | 2.91 |
| 5.0 | always vs lone | 0.569 | $3.1\times10^{-25}$ | 4.98 | 11.34 | 3.02 | 12.19 | 499 | 98 | 2.76 | 4.74 |
| 5.0 | persistent vs lone | 0.553 | $1.4\times10^{-24}$ | 6.04 | 11.34 | 6.25 | 12.19 | 615 | 98 | 2.91 | 4.74 |
| 7.0 | always vs persistent | 0.309 | $5.0\times10^{-24}$ | 4.95 | 5.86 | 3.01 | 6.66 | 499 | 634 | 2.83 | 2.16 |
| 7.0 | always vs lone | 0.500 | $2.1\times10^{-25}$ | 4.95 | 9.13 | 3.01 | 7.68 | 499 | 139 | 2.83 | 5.18 |
| 7.0 | persistent vs lone | 0.484 | $6.1\times10^{-25}$ | 5.86 | 9.13 | 6.66 | 7.68 | 634 | 139 | 2.16 | 5.18 |
| 9.0 | always vs persistent | 0.319 | $1.1\times10^{-27}$ | 5.00 | 5.81 | 3.12 | 6.47 | 499 | 760 | 2.77 | 2.13 |
| 9.0 | always vs lone | 0.381 | $2.6\times10^{-26}$ | 5.00 | 6.28 | 3.12 | 5.71 | 499 | 333 | 2.77 | 2.92 |
| 9.0 | persistent vs lone | 0.287 | $2.6\times10^{-17}$ | 5.81 | 6.28 | 6.47 | 5.71 | 760 | 333 | 2.13 | 2.92 |

**Table S3**. Overview of titrated residues in the apoferritin protomers. Residues listed were adjusted during constant-p*H* simulations. Protonation states are specified where relevant (e.g., H62 protonated on NE2, H170 protonated on ND1).

| Apoferritin protomer | Titrated residues |
|---|---|
| A | all except for E24, E59, H62 (proton on NE2), H170 (proton on ND1) |
| B | K143, D147, K154 |
| C | - |
| H | D39, D41, D42, K46, K169 |
| N | H115, D123, D128, E131 |
| P | H115, D128, E131, K143 |
| Q | D39, H57, E64, K68, D81, K83, K84 |
